# Niche-Based Priority Effects Predict Microbe Resistance to *Erwinia amylovora* in Pear Nectar

**DOI:** 10.1101/2024.07.03.601912

**Authors:** Christopher S. McDaniel, Rachel L. Vannette, Alondra Arroyo-Flores, Kyria Boundy-Mills, David W. Crowder, Michelle M. Grilley, Honey Pathak, Robert N. Schaeffer

## Abstract

Fire blight is a devastating disease affecting pome fruit trees that is caused by *Erwinia amylovora* and leads to substantial annual losses worldwide. While antibiotic-based management approaches like streptomycin can be effective, there are concerns over evolved resistance of the pathogen and non-target effects on beneficial microbes and insects. Using microbial biological control agents (mBCAs) to combat fire blight has promise, but variable performance necessitates the discovery of more effective solutions. Here we used a niche-based predictive framework to assess the strength of priority effects exerted by prospective mBCAs, and the mechanisms behind growth suppression in floral nectar. Through *in vitro* and *in vivo* assays, we show that antagonist impacts on nectar pH and sucrose concentration were the primary predictors of priority effects. Surprisingly, overlap in amino acid use, and the degree of phylogenetic relatedness between mBCA and *Erwinia* did not significantly predict pathogen suppression *in vitro*, suggesting that competition for limited shared resources played a lesser role than alterations in the chemical environment created by the initial colonizing species. We also failed to detect an association between our measures of *in vitro* and *in vivo Erwinia* suppression, suggesting other mechanisms may dictate mBCA establishment and efficacy in flowers, including priming of host defenses.

## INTRODUCTION

Globally, and for over a century, apple (*Malus x domestica*) and pear (*Pyrus communis*) production has been threatened by fire blight, a disease caused by the bacterial pathogen *Erwinia amylovora* (hereafter *Erwinia*) (van der Zwet et al. 2016). Infection by *Erwinia* occurs through the flower, with the pathogen entering hosts through specialized stomata called nectarthodes in the hypanthium (Thomson 2000; Pusey 2000). Fire blight causes wilting of shoots, necrosis of flowers, fruits, and leaves, and the formation of canker-like lesions on woody tissues that serve as overwintering inoculum (Thomson 2000). The bacterium reaches flowers through rain-, wind-, or pollinator-mediated dispersal (Johnson et al. 1993) and thrives in warm, humid conditions. *Erwinia* populations increase on the stigma before migrating through nectar into the host, a process facilitated by rain and dew (Pusey 2000). Given the losses experienced by producers, finding effective means to manage this disease are imperative (Bonn and van der Zwet 2000).

To date, producers largely rely on cultural practices and antibiotics such as streptomycin for control; however, antibiotics select for resistance in *Erwinia* populations and other bacteria in the orchard environment (Fӧrster et al 2015). Thus, antibiotics are tightly regulated in Europe and have been prohibited in US organic orchards since 2014, spurring a need for alternative management strategies (Johnson and Temple 2013; Stockwell and Duffy 2012; Fried et al. 2013). Fortunately, non-antibiotic solutions to *Erwinia* control have emerged and show promise in attenuating growth and infection of apple and pear. These strategies typically involve the use of a copper bactericide applied in late fall to limit survival of *Erwinia* in overwintering cankers, a fruit-load thinning agent at bloom to reduce potential for *Erwinia* outbreaks, a microbe treatment mid-bloom to limit epiphytic pathogen growth, and finally, full-bloom and petal-fall treatments of non-antibiotics (Johnson and Temple 2013; Johnson and Temple 2022; Elkins et al. 2015).

Amidst these strategies, microbial biological control agents (mBCAs) have garnered attention as key components of integrated pest management (IPM) approaches for fire blight. Research has focused on assessing their efficacy in suppressing *Erwinia’s* growth and virulence, exploring mechanisms such as resource competition, antibiosis, cell lysis, and induced defense (Cabrefiga et al. 2007; Pusey et al. 2008; Pusey et al. 2009; Zeng et al. 2023). Several species and strains have shown potential in reducing fire blight when applied in an IPM program (Zeng et al. 2023; Xu and Jeger 2013; Vannette and Fukami 2014; Janisiewicz and Korsten 2002). For example, BlossomProtect (*Aureobasidium pullulans*) has shown an 82% reduction in fire blight symptoms, comparable to streptomycin, leading to its widespread adoption in organic orchards across the Pacific Northwest USA (Kunz et al. 2008; Johnson et al. 2020; Zeng et al. 2023). However, *Aureobasidium* can cause fruit russeting, adversely affecting yield and marketability in wetter environments like the East Coast and Midwestern USA. (Heidenreich et al. 1997). There are additional mBCA products on the market, although they vary in efficiency and many perform inadequately relative to streptomycin (Pusey and Wend 2012; Roselló et al. 2013). Given these challenges, there is a growing need to discover novel strains with potential to suppress growth and virulence of *Erwinia*, along with an emphasis on the mechanisms underlying these processes (Pusey and Wend 2012; Cabrefiga 2011; Pusey et al. 2009; Mechan Llontop et al. 2020).

Insights from community ecology studies on the floral microenvironment hold promise for the discovery of novel mBCAs. Understanding dynamic interactions among floral microbes could help identify key characteristics shared by potent mBCAs, as flowers are highly ephemeral as compared to other host tissues and are exposed to ultraviolet light and other environmental stressors, which can impose stress on the pathogen and mBCAs applied. Furthermore, microbial establishment often depends the order of species arrival (Fukami 2015) and the microbe’s capability to tolerate additional chemical stresses imposed by the flower environment (Lievens et al. 2015; Morales-Poole et al. 2022). Floral nectar often contains high levels of sugars as a means to attract pollinators, which imposes high osmotic pressure on species attempting to colonize this resource (Lievens et al. 2015). Nectar also contains low levels nitrogen in the form of amino acids (Peay et al. 2012; Dhami et al. 2016). As such, floral microbial communities tend to be species-poor and a great competitive advantage is often conferred to the first species to arrive (Fukami 2015; Pozo et al. 2011). Commonly, the species that arrives first suppresses the growth of subsequent species by depleting key resources, or heavily modifying the nectar environment (Mittlebach et al. 2016; Peay et al. 2012; Vannette and Fukami 2014). Priority effects exerted by early arriving species can often be predicted through dissection of their niche, including overlap, impact, and resource requirements with respect to both the floral environment and later arriving species (Vannette and Fukami 2014). Here, we employ a niche-based predictive framework (Vannette and Fukami 2014) to predict the strength of priority effects exerted by prospective mBCAs against *Erwinia*. More specifically, we characterize the growth and resource use of 44 strains of flower- and insect-associated microbes in two pear nectar analogs and assess each strain’s ability to suppress *Erwinia* growth *in vitro*. Using these growth and resource-use metrics as predictors in a linear model of *Erwinia* growth suppression, we aim to understand the traits shared by species with the greatest suppressive ability. Finally, we used an *in vivo* assay to determine if our *in vitro* results are predictive of performance in actual pear flowers.

## MATERIALS AND METHODS

### Microbial strains

We assembled a diverse set of plant- and insect-associated microbes for this study (Table 1). Briefly, 44 of these strains were isolated from apple and pear flowers collected in 2017 from orchards spanning the Wenatchee River Valley (Washington), a major pome fruit producing region in the USA. Flowers with fully reflexed petals were collected at peak bloom, then returned to the lab for processing, where whole flowers were both suspended and agitated in 1× 0.15% phosphate-buffered saline-Tween solution to free epiphytic microbes. Subsamples were then dilution plated on selective media to culture fungi [yeast malt agar (YMA) + chloramphenicol (100 mg L^−1^)] and bacteria [R2A + cycloheximide (100 mg L^−1^)].

**Table 1.** (a.) Regression coefficients from the final model predicting the effects of prospective microbial (fungi and bacteria) biological control agent niche components on *Erwinia amylovora* growth. (b.) Regression coefficients from the final model predicting the effects of prospective bacterial biological control agents on *Erwinia amylovora* growth.

| Predictors | Coefficient | Standard Error | *P*-value | Reduction in $R^2$ |
| --- | --- | --- | --- | --- |
| (Intercept) | 1.49 | 0.81 | 0.028 |  |
| Total Amino Consumption | -2.10 | 0.46 | < 0.001 | 0.007 |
| pH | 1.17 | 0.18 | < 0.001 | 0.057 |
| R.PC2 | -0.70 | 0.09 | < 0.001 | 0.045 |
| Background | 1.22 | 0.2 | < 0.001 | 0.006 |
| (Intercept) | 12.27 | 2.06 | < 0.001 |  |
| Overlap | 0.03 | 0.01 | 0.0034 | 0.008 |
| Distance | -1.86 | 0.52 | 0.0017 | 0.014 |
| Background | 1.10 | 0.32 | < 0.001 | 0.018 |
| R.PC1 | -0.62 | 0.09 | < 0.001 | 0.034 |
| R.PC2 | -0.93 | 0.16 | < 0.001 | 0.068 |

Select colonies were then sub-cultured and identified using 16S and ITS sequencing. Additional strains associated with flower-visiting insects and a wildflower (*Epilobium canum*) were secured from the Phaff Yeast Culture and Vannette Lab collections (UC Davis, USA). The majority of bacterial and fungal strains were cultured on DIFCO R2A and YMA media (Becton Dickinson, Sparks, MD, USA) respectively, supplemented with 75g/L sucrose and at 27°C for three days prior to use in the growth and priority effect experiments below. *Erwinia* strains Ea8, Ea8R, and Ea8KI were grown on Luria-Bertani (LB) agar under similar conditions. Ea8 is a strain of *Erwinia amylovora* that was isolated from an infected pear tree, and Ea8R is a spontaneous mutant of strain Ea8 that is resistant to the antibiotic rifampin at concentrations up to 100mg/L. *Erwinia* strain Ea8KI, used in the flower-freezing assay below, is a transconjugant of Ea8R. The strain harbors the stable plasmid pJEL1703, which contains the *ice*C gene from *Pseudomonas syringae*, conferring high levels of ice-nucleation activity (Mercier and Lindow 1996).

### Microbial growth in nectar

Microbial growth was measured by inoculating artificial nectar with each antagonist separately. Two synthetic nectar solutions were used: a 15% w/v and a 7.5% w/v solution, designed to mimic conditions likely experienced by pear floral colonists. These nectar analogs mirrored sugar concentrations and ratios typically found in pear nectar, consisting of sucrose, fructose, and glucose mixed in a 1:2:2 ratio (Benedek et al., 2000). The nectar was also supplemented with 500 μL of BioWhittaker high non-essential amino acid mix (Lonza, Walkersville, MD, USA), as well as 1g DIFCO BACTO-Peptone (Becton Dickinson, Sparks, MD, USA) to provide microbes with amino acids; amino acids are commonly found in floral nectar and may serve as an important limiting resource for microbial growth (Peay et al. 2012; Vannette and Fukami 2014; Dhami et al. 2016). The second nectar analog, a half-dilution of the first (7.5% w/v), was chosen to simulate nectar that could result from rain events or humid conditions when *Erwinia* infection is most common (Pusey et al. 2000).

To assess growth, a 96-well plate (Costar; Corning Inc., Corning, NY, USA) was used, and each well was filled with 250 µL of artificial nectar. Each microbe was grown in triplicate for each nectar background by pipetting 50 µL of a 400 cell/µL dilution into each well. The plates were placed in either a BioTek Cytation 5 or Synergy HT plate reader (Agilent, Santa Clara, CA, USA) set at 22°C. This temperature was chosen as it was the lowest setting available for the respective plate readers. Although lower than *Erwinia*’s optimum growth temperature, 27°C (Pusey et al. 1998, 2000; Pusey and Curry 2004), it is more in range with temperatures typically experienced by *Erwinia* and prospective mBCAs during the bloom window (usually ∼14°C in the Pacific Northwest). To test for differences in the measurements between plate readers two identical plates were run simultaneously in both readers. When comparing growth and carrying capacity metrics using a paired *t*-test, no difference in overall performance was observed (*p* = 0.89). The plate was shaken every 15 min to resuspend cells that settled out of solution, and absorbance readings were taken at 600 nm to approximate cell density. The protocol ran for 72 hr and was then terminated. Absorbance readings were exported and used with the ‘SummarizeGrowthByPlate()’ function in the ‘GrowthCurver’ package (Sprouffske, 2020) in R (R Core Team, 2022) to process optical density readings and create growth profiles for each microbe. Specifically, average growth rate, carrying capacity, and lag time-related metrics for each strain in each nectar background were calculated using an average of the three replicate wells in the assay.

### Microbial impact on the nectar environment

To assess changes in nectar chemistry from microbial metabolism, we employed two methods. The concentration of sucrose, fructose, and glucose was measured using the D-Glucose Sucrose/D-Fructose/D-Glucose Assay Kit (Megazyme, Wicklow, Ireland). For this analysis, 10 μL of nectar was filtered (Nalgene Syringe Filters 0.2 μM, Nalge Company, Rochester, NY, USA) from the growth assay plates and diluted in 990 μL of sterile water. Enzymes and substrates were then added following the manufacturer’s protocol, with the assay scaled down tenfold to suit a 96-well plate. Absorbance readings were taken at 340 nm using a BioTek Cytation 5 plate reader (Agilent).

To measure amino acid concentration, we used high-performance liquid chromatography (HPLC). A Thermo-Fisher Vanquish HPLC system (Thermo Fisher Scientific Inc., Waltham, MA, USA) equipped with a Thermo UltiMate Rapid Separation Binary Pump (HPG-3400RS; Thermo Scientific), a Thermo UltiMate autosampler (WPS-3000; Thermo Scientific), and a Thermo UltiMate column compartment (TCC-3100; Thermo Scientific) was used. For each filtered nectar sample, 50 μL was loaded into individual HPLC vials. Amino acids were derivatized *in-situ* using ortho-phthalaldehyde (Phtlalaldehyde reagent complete solution; Sigma Aldrich, Darmstadt, Germany) for primary amino acids and 9-fluorenylmethyl chloroformate (LiChropur FMOC chloride; Sigma Aldrich) for secondary amino acids. Each sample was supplemented with 1 μL of 10 Alpha-Amino-n-butyric acid (AABA; Aldrich Chemical Company, Milwaukee, WI, USA) as an internal standard. A solvent gradient was employed to separate amino acids using a borate buffer (1 L HPLC grade water, 10 mM Na2HPO4, 10 mM Na2B4O7, 0.5 mM NaN3, pH 8.2) as the polar phase and acetonitrile–methanol–water [45%, 45%, 10% (v/v), Acetonitrile; LiChrosolv Sigma Aldrich, Methanol; Fisher Chemical Optima LC/MS Fair Lawn NJ, water; Sigma Aldrich HPLC grade water] as the non-polar phase. Detection was performed at 338 nm and 262 nm for the OPA- and FMOC-derivatives, respectively, using a Thermo Vanquish Diode Array Detector (VF-D11-A; Thermo Scientific).

Finally, a pH reading was collected for each well using a Fisher accumet excel XL 25 dual channel pH/ion meter (13-636-XL25; Fisher Scientific, Waltham, MA, USA) and a Fisher accumet microglass, mercury-free, combination electrode (13-620-851; Fisher Scientific).

### Priority effects experiment

To assess the relative strength of priority effects exerted by prospective mBCAs on *Erwinia* growth, we performed an *in vitro* competitive assay, again using 96-well plates (Costar). Plates were inoculated with 90 μL of each pear nectar analog with 5 μL of a 400 cell/uL suspension of each microbe. There were 5 replicates per nectar background for each microbe. These were allowed to grow for 48 hr at 22°C in the Synergy HT plate reader (Agilent) before 5 μL of a 400 cell/uL suspension of *Erwinia* was added to each well to test the suppressive ability of each. After 120 hr, the experiment was terminated and growth of *Erwinia* was assessed by diluting each well 1000-fold, then plating a 20 μL subsample on R2A media supplemented with 75g/L sucrose; 100mg/L of either cycloheximide or rifampin were added to the media to suppress growth of yeasts and bacterium, respectively. We used two controls, the positive control being *Erwinia* grown in nectar alone for 72 hours, where the negative control used sterile nectar solution. We then assessed suppressive ability of each microbe by counting colonies and estimating colony forming units (CFUs) of *Erwinia* in each well.

### Antagonism assay in flowers

To test whether microbes suppressed growth of *Erwinia* on pear flowers, we adopted a flower-freezing assay developed by Mercier and Lindow (1996). Briefly, dormant branches were harvested from Bartlett pear trees grown in a research orchard near Davis, CA, USA. Cut branches were ∼50 cm long, containing 5-10 clusters of flower buds each. These were stored dry in a cold room (∼4°C) at the UC Davis until use. After they were brought out of storage, basal ends were excised under water, then branches were placed in containers of water to a depth of ∼5 cm. Branches were then forced to bloom by incubating them on growing shelves at ∼25°C under continuous light. Before use, branches with newly opened flowers (1-3 d old) were cut to a length of 15–25 cm, with all unopened flowers removed.

Bacterial and fungal strains to be tested were plated on R2A and YM media, respectively, and incubated at 27°C for ∼3 d before use. Bacterial and fungal cells were removed from agar with sterile toothpicks and suspended in 5 mL of 5 mM phosphate buffer, pH 7.0, to yield a suspension of ∼10^7^ cells per ml. Recently opened flowers were inoculated with strains by misting them with a spray bottle containing the cell suspension. From 2 to 4 branches with a total number of 25 to 40 flowers were inoculated with a given antagonist. Branches inoculated were enclosed in a plastic bag and incubated at room temperature (−21°C). There were two controls for each experiment: the first was composed of flowers sprayed with buffer only, and after 48 hr, the blossoms (including controls) were sprayed with a suspension of Ea8KI (∼10^4^ cells per ml) in 5 mM phosphate buffer, and the second were flowers dipped in sterilized buffer only. The branches again were enclosed in plastic bags and incubated for 48 hr at room temperature. After this incubation period, flowers were aseptically removed from treated branches and placed individually in 10 mL tubes containing 3.5 mL of 10 mM phosphate buffer. The threshold freezing temperature (i.e., the warmest temperature that caused freezing) of each flower was determined by immersing the tubes in a circulating glycerol bath at −2°C and lowering the temperature by an increment of 0.5°C every 15 min. The number of frozen tubes for each treatment was counted when 85 to 100% (typically 95%) of the control tubes (containing flowers inoculated only with Ea8KI) had frozen. The effectiveness of each strain was expressed as the relative freezing potential, which was calculated by dividing the number of flowers treated with an antagonist that had frozen by the number of flowers treated only with Ea8KI that had frozen.

### Statistical analyses

Following a modified approach from Vannette and Fukami (2014), we used a principle component regression model to examine whether the niche components measured could explain the strength of priority effects exerted by prospective mBCAs on *Erwinia*. Since antagonist growth and sugar use metrics were highly correlated with each other (Tables S1 and S3), we used principal component analyses to eliminate the correlation between variables (Tables S2 and S4). We then used the first two principal components as predictor variables in our model, which for growth and sugar use explained 81% and 93% of the variance in the data respectively (Tables S2 and S4). We started with the full linear model:

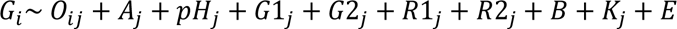

Where *G_i_* is the growth of species *i* measured in CFUs (+1, log10-transformed); *O_ij_* is the scalar product of amino acid use overlap between each microbe following (Pianka 1973) and *Erwinia*; *pH_j_* is the pH of the nectar inoculated with species *j*; *G1_j_* and *G2_j_* are the first and second principle component axes that represent growth metrics of species *j*; *R1_j_* and *R2_j_* are the first and second principle component axes that represent per capita sugar consumption of species j; B is nectar background in which species i and species j were competing; *K* is the kingdom of species *j*, and *E* is the irreducible error term. Species *i* represents *Erwinia,* and species *j* represents the microbes tested against *Erwinia.* Only Growth PC1 and PC2 had variance inflation factor (VIF) scores above 5, so we only retained PC1 in the initial full model. All other predictors were not correlated (VIF < 5), and therefore were included in the initial full model.

In a subsequent analysis focusing on bacteria only, we included a term for phylogenetic relatedness. Prior work on nectar-inhabiting microbes (Peay et al. 2012) showed competition between congener species can be more intense than between distantly related species, in line with Darwin’s naturalization hypothesis (Darwin 1859). To measure phylogenetic relatedness, the 16S sequences for bacteria were first aligned using the msa package (Bodenhofer et al 2015). The *phangorn* package (Schliep 2011) was then used to first construct a neighbor-joining tree, following by fitting of a GTR+G+I (Generalized time-reversible with Gamma rate variation) maximum likelihood tree, using the neighbor-joining tree as a starting point. We then calculated the patristic distance between all bacterial species and *Erwinia* as a predictor.

The best predictors for each model were selected using best subset regression (Hebbali 2020), comparing Akaike’s information criterion (AIC) and selecting parsimonious models within ΔAIC of 6 from the model with the lowest AIC value. We chose this higher ΔAIC (ΔAIC ≤ 2 is standard; Burnham and Anderson 2022), as it is likely required to give a 95% probability of including the best (expected) Kullback–Leibler Distance model in the top model set (Richards 2005; 2008; Harrison et al. 2018). Changes to adjusted *R^2^* for each predictor were noted, allowing us to determine the relative importance of each predictor in explaining the variance in priority effects observed in the final models (Tables 3 and 5).

Finally, for the flower freezing assay, we conducted a Spearman rank correlation (Figure 4) between *Erwinia* growth measured in the *in vitro* experiment and the proportion of flowers remaining unfrozen at −4.5°C. At this temperature setpoint, all flowers treated with *Erwinia* alone had completely frozen (i.e., no “survival”).

## RESULTS

### In vitro suppression of *Erwinia* growth

Microbes displayed considerable range in their effectiveness in suppressing *Erwinia* growth *in vitro* (Figure 1). Among those evaluated, five species were able to completely suppress *Erwinia*. Most suppression of growth occurred in the diluted pear nectar analog, however, *S. bombicola* was able to completely suppress growth in the 15% analog. Of these five species, three were yeasts (*Starmerella bombi, S. bombicola,* and *Metschnikowia mogii*), and two bacteria, *Rosenbergiella* sp. and *Micrococcus yunnannensis*.

**Figure 1.**
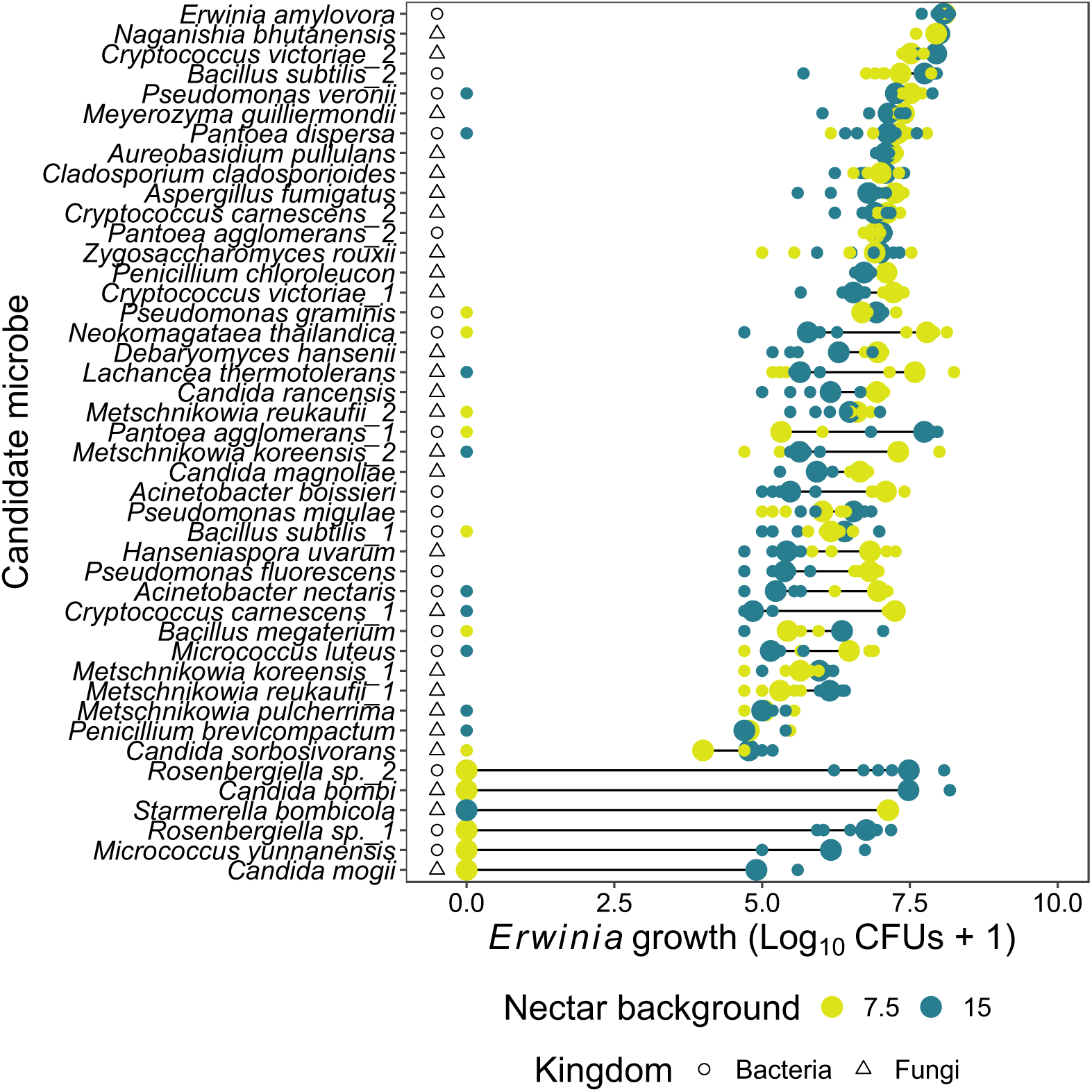
Cleveland plot depicting the impact of candidate bacteria and fungi on *Erwinia amylovora* growth [Log_10_(CFUs +1)] in two pear nectar analogs (7.5% and 15% weight by volume). Each open data point represents the individual CFU counts for that respective replicate, depicted as open points, while the larger, closed points indicate the CFU counts averaged across all five replicates. The *Erwinia amylovora* data points correspond to an assay without the presence of an antagonist (control).

**Figure 2.**
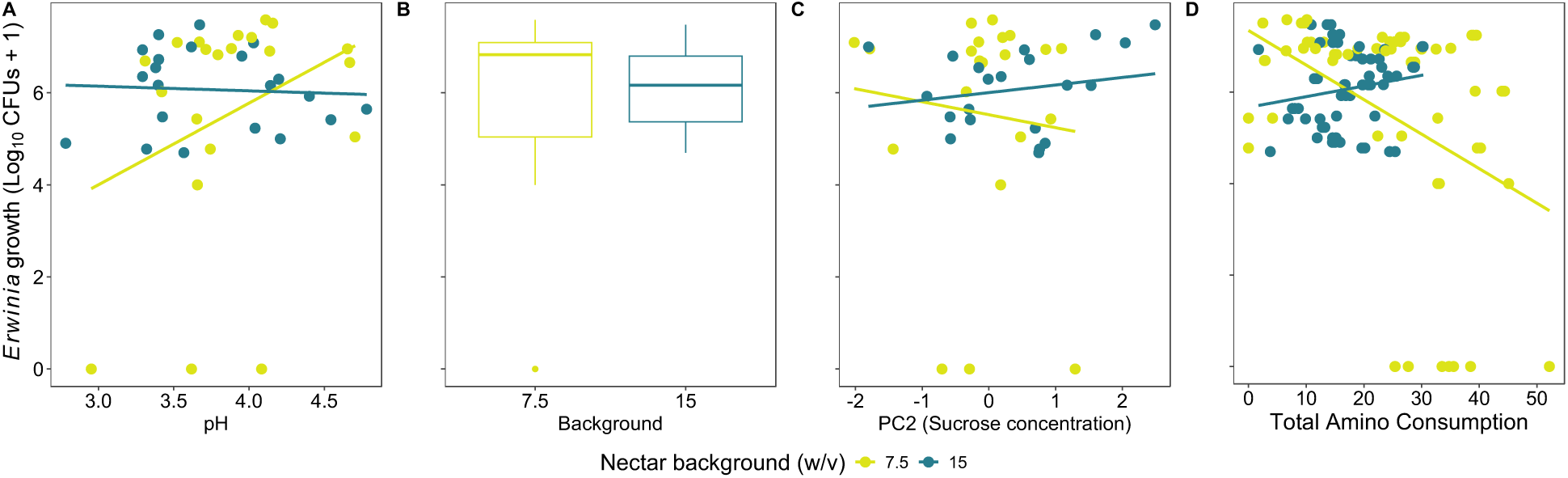
Relationships among pear nectar traits impacted by candidate microbes *in vitro* and *Erwinia amylovora* growth [Log_10_(CFUs +1)].

### Predictors of in vitro suppression

Following analysis of the dataset and full model, we retained four predictors: total amino acid consumption, pH, resource use PC2 (associated with sucrose metabolism), and nectar background (Table 1a). The overall model explained 12.3% of variance in *Erwinia* growth, with 10.1% of that attributable to shifts in nectar pH and sucrose concentration (Table 1a.). For the bacteria-only model, we retained five predictors: niche overlap, phylogenetic distance, resource use PC1 and PC2, as well as nectar background (Table 1b.). The strongest predictors were the two PCs relating to sugar concentrations, which explained 10% of the variance in *Erwinia* growth outcomes (Table 1b.).

The best predictor of *Erwinia* growth suppression in both models was per capita changes to sugar concentrations within nectar. In addition, decreases in sucrose concentration correlate with increases in both glucose (*r* = −0.33, *p* < 0.001) and fructose concentration, (*r* = −0.27, *p* < 0.001), suggesting that the microbes were cleaving sucrose into its monomers, but not decreasing the concentration of either monosaccharide drastically. Each species varied in its metabolism of sucrose, and species accounted for 57% of the variation in sucrose concentration in our data.

Species ranged considerably in their effects on nectar pH, with *Candida mogii* having the largest impact, reducing pH below 3 in both nectar backgrounds. Species explained 87% of the variance in pH values recorded. Moreover, pH was retained as a predictor in the overall model and accounted for 5.6% of the variance in *Erwinia* growth performance.

There were four strains with a high degree of niche overlap (Pianka index > 0.98). Of these, only one was closely related to *Erwinia*, *Pantoea agglomerans*, with the other three being more distantly related: *Bacillus megaterium, B. subtilis*, and *Penicillium chloroleucon*. Among the amino acids that we were able to detect via HPLC, we identified eight that were strongly associated with our overlap metric: asparagine, isoleucine, leucine, serine, threonine, glycine, phenylalanine, and valine (co-eluted with methionine). Despite this, niche overlap was a weak predictor, being retained only in the bacterial model and explaining < 1% of the variance in *Erwinia* growth performance (Table 1b).

Phylogenetic relatedness was retained as a predictor in the best overall bacterial model, however, it only explained 1.4% of the variance in the dataset (Table 1b). In our study we found that phylogenetic relatedness was weakly correlated with niche overlap using Pearson correlation (r = −0.19, *p* = <0.001); however, this does suggest that on average more closely related species have a higher degree of niche overlap.

### In vivo suppression of *Erwinia* growth

As in the *in vitro* assay, microbes evaluated in this experiment displayed extensive range in their ability to suppress *Erwinia* growth *in vivo*. Antagonistic strains [ratio of frozen flowers (treated/control): 0-0.3] included yeasts *Candida rancensis*, *M. koreensis* (strain 1), *Zygosaccharomyces rouxii* and bacteria *Micrococcus* spp., *P. graminis*, and *Rosenbergiella* spp. (Figure 3). A Spearman rank correlation test revealed no significant association in *Erwinia* growth outcomes between the *in vitro* and *in vivo* assays however (*r* = −0.24, *p* = 0.17, Figure 3).

**Figure 3.**
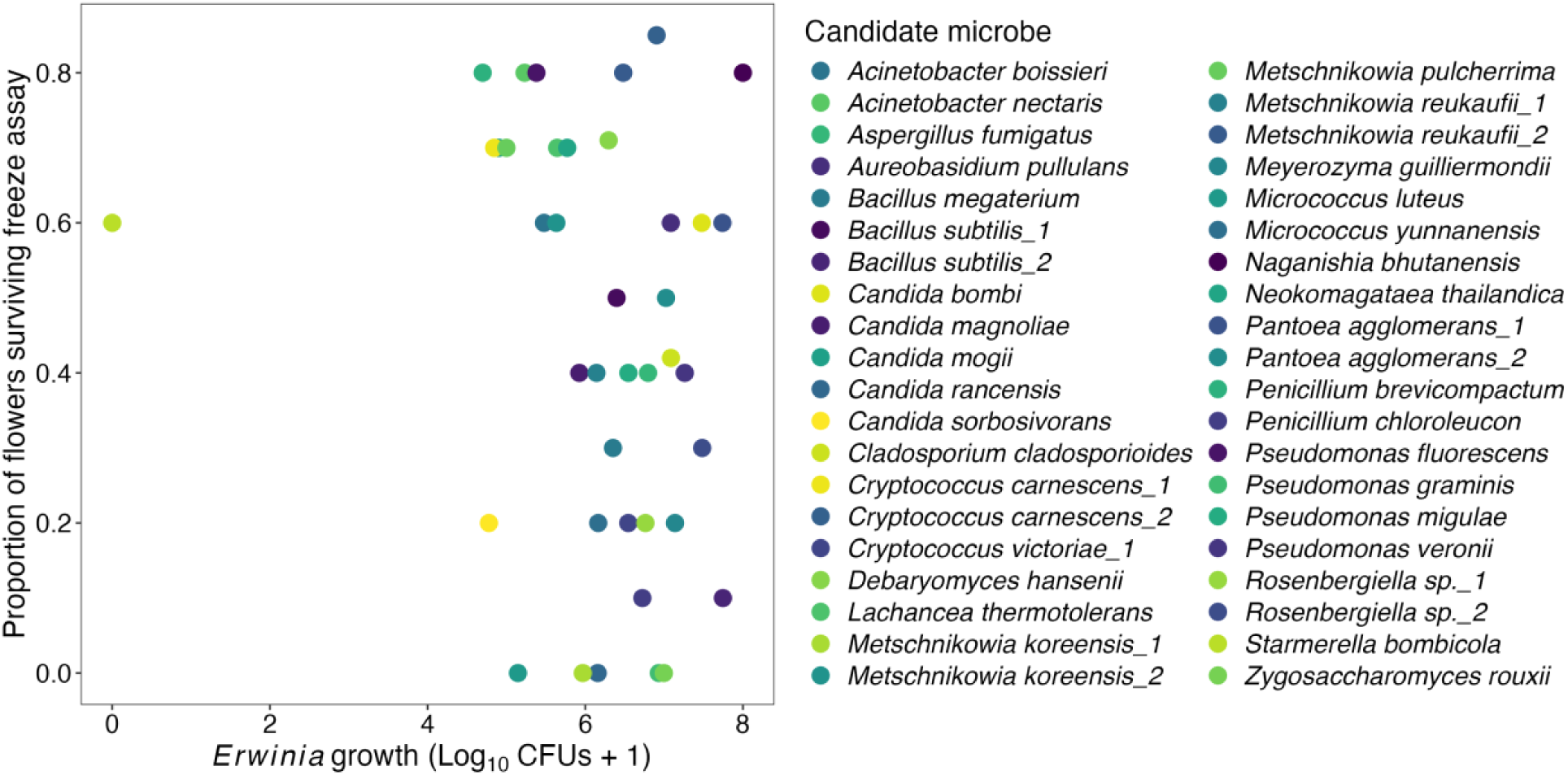
Association (Spearman rank correlation) between results of the *in vitro* priority effects assay antagonist suppression of *Erwinia amylovora* growth *in vitro* [Log_10_(CFUs +1)] and *in vivo* (proportion surviving during the flower freezing assay) where greater survival (higher proportion) indicates a lower abundance of *Erwinia*.

## DISCUSSION

Antagonists tested in this study displayed considerable range in their ability to suppress *Erwinia* growth both *in vitro* and *in vivo*. Of the 40+ strains tested, only a handful suppressed the pathogen fully, and largely only in the 7.5% pear nectar analog. The microorganisms that suppressed growth the most were those that drastically reduced nectar pH, in line with work that noted an important role of pH in affecting *Erwinia* growth and virulence (Wodzinski et al. 1994; Pester et al 2012), as well as perhaps a more general pattern of pH in mediating priority effects exerted among microbes during community succession (Chappell et al. 2022; Fierer and Jackson 2006). Previous research on *Erwinia* has shown that its growth is hindered at pH levels below 5 (Raymundo 1980; Pusey et al. 2008), and viability further at levels below 4 (Wodzinski et al. 1994). *Erwinia*’s optimal growth conditions are between 20-28°C and pH r 6-8 (Raymundo et al. 1980). At pH levels of 4 and below *Erwinia* loses a key virulence factor, its type III secretion system, which aids in suppression of the host’s immune response; thus, making it less effective at infection (Pester et al 2012). More generally, a recent study on nectar microbes corroborates our result, showing that priority effects in monkeyflower (*Diplacus auranticus*) nectar are mainly governed by reductions in pH from the early arriving species (Chappell et al. 2022). A disclaimer to note regarding our experimental approach is that pH readings were taken 72 hr after initiation of our single strain growth assays. In our priority effects experiment, *Erwinia* was introduced to nectar environments 48 hr after the start. It is very likely that the nectar we introduced *Erwinia* into in the priority effects experiment had slightly higher pH levels than those used as predictors in our model. This would explain the apparent, albeit limited, growth of *Erwinia* in nectar with a pH below 4. Another caveat to note is that we plated *Erwinia* on non-acidic media following the competition assay, therefore, even if only a few cells survived, it is likely that they would be able to form colonies when introduced to the fresh media. reduced nectar pH, with most belonging to closely related nectar-inhabiting yeasts.

Another factor associated with growth suppression was related to the ratio of sugars in the nectar environment following antagonist growth and metabolism. Microbes that metabolized sucrose, leading to a proportional increase in nectar monosaccharides (glucose and fructose), showed the poorest performance in the suppression trial. This suggests that the harshness of the nectar environment in our study was not solely driven by sugar concentration but rather by the mono- to di-saccharide ratio. Solutions rich in monosaccharides have higher molarity, leading to lower water activity and stronger osmotic pressure even at equivalent % w/v (Velezmoro, Oliveira and Meirelles 2000). This may have an inhibitory effect even on microbes that tolerate high osmotic pressures well, such as nectar microbes (Morales-Poole et al. 2022), however this does not seem to be the case for *Erwinia.* Previous research has shown that *Erwinia* outgrows other nectar microbes (*Pseudomonas fluorescens* and *Pantoea agglomerans)* in nectar at high mono- to di-saccharide ratios (2:1, 1:0) (Pusey 1998). Moreover, a study using similar nectar microbe species to those used in our study (*Acinetobacter* and *Rosenbergiella* spp.) showed that in 50% w/v artificial nectar growth was only supported in the solutions comprised entirely of sucrose, and nectars made from monosaccharides inhibited growth entirely (Morales-Poole et al 2022). Our synthetic nectar analog began at a 2:1 mono- to di-saccharide ratio, and with the high metabolism of sucrose by most microorganisms tested, became more hexose dominant over time. This would explain the low levels of growth suppression observed, as *Erwinia* was introduced to a nectar environment with higher osmotic pressure, as it tolerates those conditions better than most microbes (Pusey 1998). The ability of *Erwinia* to tolerate nectars with high monosaccharide content provides it with a competitive advantage over most nectar microbes tested, making tolerance to these conditions a critical trait to consider when screening for novel mBCAs.

In both models, measures of amino acid consumption (total and overlap) were retained as significant predictors. We hypothesize that some of the *Erwinia* suppression observed can be attributable to antagonist depletion of specific amino acids, which are a limiting resource for microbes that colonize the floral nectar environment (Peay et al. 2012; Vannette and Fukami 2014; Morales-Poole et al. 2022). More specifically, we identified eight amino acids associated with a high score for niche overlap; however, the amino acids that we identified are all important for microbial metabolism and can be readily used via the Krebs cycle. For instance, one study has revealed that consumption of asparagine by *Erwinia* is common, as it is one of the main nitrogen transport molecules in xylem of apple trees (Klee et al. 2019). It is also interesting to note that in our study, we observed a very weak correlation between niche overlap and phylogenetic distance, suggesting that the metabolic profile of each microorganism cannot be attributed to its shared evolutionary history. However, we only tested a small subset of potential amino acids that can be found in floral nectar, and it is possible that we did not capture the full breadth of amino acid requirements for each species. While saying that, amino acids have been shown to be one of the main drivers of priority effects in floral nectar, due to their low concentrations and high nutritional value (Peay et al. 2011; Dhami et al. 2016). In fact, a similarly designed competition study, comparing phylogenetically-related yeasts revealed that priority effects between species are a result of high niche overlap and high rates of resource scavenging (Dhami et al. 2016). Moreover, the strong priority effects were ameliorated after addition of an amino acid mixture to the nectar environment (Dhami et al. 2016). Similarly, it has been documented that nectar microbial communities reach higher densities when pollen is added into floral nectar, due increased amino acids resulting from digestion of the pollenkitt (Herrera et al. 2010; Rering et al. 2021). This can be coupled with the fact that certain nectar specialist microbes have specific adaptations to burst pollen grains (Christenson et al. 2021). Finally, work on *Erwinia* and its nutritional requirements show an important role for specific amino acids such as arginine in virulence metabolic pathways and growth (Klee et al. 2019).

Additional direct (e.g., antibiosis; Wodzinski et al. 1994) and indirect (e.g., host defense priming) mechanisms of microbial biocontrol not measured here may be important in explaining *Erwinia* performance. With respect to the latter, *Aureobasidium pullulans* (BlossomProtect) was ranked moderate in its ability to suppress *Erwinia* growth *in vivo* and *in vitro*, mirroring previous work (Johnson 2022, Slack 2019). Despite its limited ability to curb *Erwinia* growth, a recent finding by Zeng et al. (2023) demonstrated the mode of action through which *Aureobasidium* affects this bacterial pathogen and its ability to infect pome hosts. *Aureobasidium*, when applied during bloom, activates the plants SAR response, which primarily upregulates transcription of a pair of pathogenesis related genes (*PR1* and *PR2*) that prime the plant immune response, leading to synthesis of ꞵ-1,3-Glucanases responsible for degrading the cell wall of invading pathogens (Zeng et al. 2023; Perrot et al. 2022). Moreover, *Erwinia* is able to gain entry to the plant by transiently suppressing the SAR system in the hypanthium through its type III secretion system (Cui et al. 2021). Despite the suppression of SAR by *Erwinia,* BlossomProtect restores SAR activation in the presence of *Erwinia*, protecting the plant from infection (Pester 2012; Zeng et al. 2023). Results from our study (lack of correlation between *in vitro* and *in vivo* suppression) and Zeng’s (2023) point to a role for host defense priming in limiting *Erwinia* infection. Finally, two drawbacks to note regarding BlossomProtect are: (1) the pathogenicity of *Aureobasidium* itself, as produces a suite of enzymes (e.g., cutinases and lipases) that cause fruit russeting (Johnson et al. 2022; Matteson-Heidenreich et al. 1997), affecting fresh fruit marketability and financial returns (Spotts and Cervantes 2007; Winkler et al 2022), and (2) its potential negative effect on the attraction of floral visitors, as studies suggest that it may be less attractive than other floral microbes (Rering et al. 2018; Colda et al. 2021; Crowley-Gall et al. 2022).

Further gains in microbial-based solutions for disease control could occur through development of multi-strain consortia, which may outperform single strain treatments (Niu et al. 2020). These consortia may consist of microbial species with complementary modes of action or a mix of representative strains from the same species, exhibiting effectiveness under different environmental conditions (Niu et al. 2020). The principle of functional diversity, wherein microbes are paired based on having complementary suppressive mechanisms may confer greater inhibition of growth and virulence against a pathogen (Christenson and Yannarell 2022; Carmona et al. 2016). For instance, combining *Aureobasidium* with another strain capable of suppressing *Erwinia* growth at the stigma, like *Metscnhikowia mogii,* could enhance the efficacy of such treatments (Christenson and Yannarell 2022). Additionally, considering the impact of high nectar acidity on *Erwinia’s* ability to repress the plants systemic acquired resistance (SAR) response, it might be advantageous to pair acid-producing microbes like *Metschnikowia reukaufii* with acid-tolerant fungi like *Aureobasidium* in a multi-strain biocontrol approach (Saur et al. 2019; Pester et al. 2012; Tucker and Fukami 2014).

In conclusion, we show that priority effects exerted by microbial antagonists on *Erwinia* growth *in vitro* was mainly attributable to antagonist-driven changes in nectar pH and sugar ratios. A lack of association between these results and those from the *in vivo* assay, however, combined with more recent work (Zeng et al. 2023), suggests that host immune priming may be another mechanism to exploit for microbial-mediated biocontrol of this disease in pome fruit hosts. Nonetheless, microbes that dramatically reduce nectar pH, and those that are tolerant of high monosaccharide nectars, may still confer some benefit to growers looking for more robust biocontrol. In the future, multi-strain consortia may have the potential to improve upon existing treatment options (Burgess and Schaeffer 2022) and warrants further investigation for improving both disease management and yield outcomes in an economically-important fruit, pear.

## Acknowledgments

We thank A. Crowley-Gall, C. Davis, J. Duque, G. Hall, and A. Khan for laboratory assistance, and K. Beard, M. Borghi, and V. Martin for comments on an earlier version of this manuscript.

---

**Table S.1.** Microbial strains used in this study, along with their isolate source.

| Species | Kingdom | Strain ID(s) | Isolation source |
| --- | --- | --- | --- |
| *Acinetobacter boissieri* | Bacteria | EC_115 | *Epilobium canum* |
| *Acinetobacter nectaris* | Bacteria | EC_033 | *Epilobium canum* |
| *Aspergillus fumigatus* | Fungi | EC_088 | *Epilobium canum* |
| *Aureobasidium pullulans* | Fungi | Gall_For_Edge_N_Y1 | *Prunus dulcis* |
| *Bacillus megaterium* (syn: *Preista megaterium*) | Bacteria | *B. meg* |  |
| *Bacillus subtilis* | Bacteria | EC_116, *B. sub* | *Epilobium canum* |
| *Candida bombi* (syn: *Starmerella bombi*) | Fungi | UCDFST 02-329 | *Apis mellifera* |
| *Candida magnoliae* (syn: *Starmerella magnoliae*) | Fungi | UCDFST 02-257 | *Apis mellifera* |
| *Candida mogii* (syn: *Metschnikowia mogii*) | Fungi | UCDFST 67-79 | *Bombus* sp. |
| *Candida rancensis* (syn: *Metschnikowia mogii*) | Fungi | Baugh_Org_Int_N_Y1 | *Apis mellifera* |
| *Candida sorbosivorans* (syn: *Starmerella sorbosivorans*) | Fungi | EC_094 | *Epilobium canum* |
| *Cladosporium cladosporioides* | Fungi | EC_089 | *Epilobium canum* |
| *Cryptococcus carnescens* (syn: *Vishnacozya carnescens*) | Fungi | EC_072, Baugh_Org_Int_N_Y1 | *Epilobium canum*, *Pi* |
| *Cryptococcus victoriae* (syn: *Vishnacozya victoriae*) | Fungi | Muller_Con_Int_N_Y1, Roll_Baugh_Org_Int_Y1 | *Prunus dulcis* |
| *Debaryomyces hansenii* | Fungi | UCDFST 67-77 | *Bombus* sp. |
| *Hanseniaspora uvarum* | Fungi | Asa_Int_AS_Y1 | *Prunus dulcis* |
| *Lachancea thermotolerans* | Fungi | UCDFST 68-140 | Apidae |
| *Metschnikowia koreensis* | Fungi | EC_061, EC_064 | *Epilobium canum* |
| *Metschnikowia pulcherrima* | Fungi | Asa_Edge_AS_Y2 | *Prunus dulcis* |
| *Metschnikowia reukauffii* | Fungi | EC_052, EC_067 | *Epilobium canum* |
| *Myerozyma guilliermondii* | Fungi | UCDFST 02-286 | Apidae |
| *Micrococcus luteus* | Bacteria | EC_006 | *Epilobium canum* |
| *Micrococcus yunnanensis* | Bacteria | EC_005 | *Epilobium canum* |
| *Naganishia bhutanensis* | Fungi | UCDFST 10-272 | Megachilidae |
| *Neokomagataea thailandica* | Bacteria | EC_112 | *Epilobium canum* |
| *Pantoea agglomerans* | Bacteria | Champs_P_B2, MAV_P_B2 | *Malus x domestica* |
| *Pantoea dispersa* | Bacteria | P8_P_B2 | *Pyrus communis* |
| *Penicillium brevicompactum* | Fungus | EC_137 | *Epilobium canum* |
| *Penicillium chloroleucon* | Fungus | EC_84 | *Epilobium canum* |
| *Pseudomonas fluorescens* | Fungus | P8_P_B3 | *Pyrus communis* |
| *Pseudomonas graminis* | Bacteria | P2_P_B2 | *Pyrus communis* |
| *Pseudomonas migulae* | Bacteria | P8_P_B1 | *Pyrus communis* |
| *Pseudomonas veronii* | Bacteria | MAV_P_B1 | *Malus x domestica* |
| *Rosenbergiella* sp. | Bacteria | EC_124 | *Epilobium canum* |
| *Rosenbergiella* sp. | Bacteria | EC_126 | *Epilobium canum* |
| *Starmerella bombicola* | Bacteria | UCDFST 02-305 | *Apis mellifera* |
| *Zygosaccharomyces rouxii* | Fungi | UCDFST 68-105 | *Apis mellifera* |

**Table S.2.**

| Models | Model terms |  |  |
| --- | --- | --- | --- |
| 1 | pH |  |  |
| 2 | Total amino consumption + pH + Sugar use PC2 + Nectar background |  |  |
| 3 | Total amino consumption + pH + Sugar use PC1 & PC2 + Nectar background |  |  |
| 4 | Total Amino Consumption + pH + Growth PC1 + Sugar use PC1 & PC2 + kingdom + background |  |  |
| 5 | Niche overlap + Total Amino Consumption + pH + Growth PC1 + Sugar use PC1 & PC2 + kingdom + background |  |  |
| Model | ΔAIC | AIC | Terms in model |
| 2 | 0.00 | 5874.97 | Total amino consumption + pH + Sugar use PC2 + Nectar background |
| 3 | 0.39 | 5875.36 | Total amino consumption + pH + Sugar use PC1 & PC2 + Nectar background |
| 4 | 1.60 | 5876.57 | Total Amino Consumption + pH + Growth PC1 + Sugar use PC1 & PC2 + kingdom + background |
| 5 | 2.30 | 5877.27 | Niche overlap + Total Amino Consumption + pH + Growth PC1 + Sugar use PC1 & PC2 + kingdom + background |
| 1 | 66.82 | 5941.79 | pH |

**Table S.3.**

| Models | Model terms |  |  |
| --- | --- | --- | --- |
| 1 | Sugar use PC2 |  |  |
| 2 | Sugar use PC1 & PC2 |  |  |
| 3 | Sugar use PC1 & PC2 + Nectar background |  |  |
| 4 | Niche overlap + Phylogenetic distance + Sugar use PC1 & PC2 + Nectar Background |  |  |
| 5 | Niche overlap + Phylogenetic distance + Growth PC1 + Sugar use PC1 & PC2 + Nectar Background |  |  |
| Model | ΔAIC | AIC | Terms in model |
| 5 | 0 | 2234.66 | Niche overlap + Phylogenetic distance + Growth PC1 + Sugar use PC1 & PC2 + Nectar Background |
| 4 | 1.91 | 2236.57 | Niche overlap + Phylogenetic distance + Sugar use PC1 & PC2 + Nectar Background |
| 3 | 9.55 | 2244.21 | Sugar use PC1 & PC2 + Nectar background |
| 2 | 13.4 | 2248.04 | Sugar use PC1 & PC2 |
| 1 | 40.8 | 2275.49 | Sugar use PC1 |

